# Ribo-seq guided design of enhanced protein secretion in *Komagataella phaffii*

**DOI:** 10.1101/2025.02.12.637931

**Authors:** Aida Tafrishi, Troy Alva, Justin Chartron, Ian Wheeldon

## Abstract

The production of recombinant proteins requires the precise coordination of various biological processes, including protein synthesis, folding, trafficking, and secretion. The overproduction of a heterologous protein can impose various bottlenecks on these networks. Identifying and alleviating these bottlenecks can guide strain engineering efforts to enhance protein production. The methylotrophic yeast *Komagataella phaffii* is used for its high capacity to produce recombinant proteins. Here, we use ribosome profiling to identify bottlenecks in protein secretion during heterologous expression of human serum albumin (HSA). Validation of this analysis showed that the knockout of non-essential genes whose gene products target the ER, through co- and post-translational mechanisms, and have high ribosome utilization can increase production of a heterologous protein, HSA. A triple knockout in co-translationally translocated carbohydrate and acetate transporter Gal2p, cell wall maintenance protein Ydr134cp, and the post-translationally translocated cell wall protein *Aoa65896.1* increased HSA production by 35%. This data-driven strain engineering approach uses cell-level information to identify gene targets for phenotype improvement. This specific case identifies hits and creates strains with improved HSA production, with Ribo-seq and bioinformatic analysis to identify non-essential ER targeted proteins that are high ribosome utilizers.

## 1. Introduction

Engineering diverse biologics such as enzymes, antibodies, protein therapeutics, and materials is a large part of industrial biotechnology and biopharma (Sanchez-Garcia et al., 2016). *Komagataella phaffii* (*K. phaffii*), commonly known as the ‘biotech yeast’(Bernauer et al., 2020), stands out among the fungal kingdom for its ability to express and secrete proteins. This yeast grows to high cell densities, secretes low levels of endogenous proteins simplifying downstream processes, has a native promoter system that is tightly regulated under methanol condition, and induces high expression levels thus enabling heterologous protein production (Bernauer et al., 2020). Protein overexpression in *K. phaffii* typically involves growing cells in glycerol-based media to produce biomass followed by switching to methanol-based media for heterologous induction, using the methanol inducible *AOX1* promoter for expression (Cereghino and Cregg, 2000; Gasser et al., 2006; Liang et al., 2012).

Various strategies have been employed to increase heterologous production including optimization of growth conditions, engineering signal sequences, and refactoring secretory pathway expression (Bae et al., 2015; de Ruijter et al., 2016; Gustafsson et al., 2004; Huang et al., 2018; Karbalaei et al., 2020; Mori et al., 2015; Narendranath and Power, 2005; Payne et al., 2008; Salari and Salari, 2017). These approaches are effective but many bottlenecks in the secretory pathway are unknown and remain unaddressed (Mattanovich et al., 2004; Wang et al., 2017). A known rate-limiting step in production is protein trafficking through the endoplasmic reticulum (ER) (Zahrl et al., 2018). Heterologous trafficking relies on the recognition and binding of N-terminus hydrophobic signal sequences by the cell’s translocation machinery (Akopian et al., 2013; Zahrl et al., 2018). Signal Recognition Particles (SRPs) and other proteins in the secretory pathway guide ribosome nascent chain (RNC) complexes to the ER membrane where they interact with co- and post-translational translocons (Nyathi et al., 2013). A deficiency of chaperones can lead to an unfolded protein response or intracellular retention of misfolded/unglycosylated proteins. Access to protein folding chaperones in *K. phaffii* is made more difficult as an equal amount of nascent polypeptides translocate across the ER co- and post-translationally (Alva et al., 2021).

Ribosome profiling (Ribo-seq) is a next-generation sequencing technique that measures protein synthesis by assessing ribosome abundance at each codon in a transcript (Ingolia, 2016). Compared to RNA-seq, Ribo-seq offers a closer correlation with standard proteomics and is much higher throughput than mass spectrometry, while still accurately predicting mature protein stoichiometry (de Bruijn, 2016; Li et al., 2014; Taggart and Li, 2018). Unlike mass spectrometry, which is influenced by degradation, Ribo-seq specifically reports on protein synthesis, providing a more direct measurement of translational activity. However, Ribo-seq comes with technical challenges, particularly in the isolation of ribosome-protected mRNA footprints, which requires effective rRNA subtraction methods (Archer et al., 2014; Chung et al., 2015; Faridani et al., 2016; Ingolia et al., 2012).

Here, we take a data-driven strain engineering approach that uses cell-level information on ribosome utilization to guide strain engineering of *K. phaffii* GS115 to improve heterologous protein production. We first conducted a Ribo-seq screen to help understand the protein expression landscape in *K. phaffii*, focusing on protein translocation and secretion. We explored methanol metabolism and employed Ribo-seq alongside ER trafficking predictions to generate data reflecting translatome variations in wildtype and human serum albumin (HSA)-expressing strains. After creating a cellular map of protein expression with Ribo-seq, we used this information to inform our strain engineering. Multigene knockout of key, non-essential host cell proteins involved in early secretory pathways resulted in significantly improved HSA secretion. Our methodology reveals novel insights into these conditions and allows for a rational approach to widen secretion bottlenecks by providing new targets for modification that would not have otherwise been predicted.

## 2. Results

### 2.1 Surveying translation with Ribo-seq

We used a high-throughput Ribo-seq technique to measure protein synthesis for wildtype (Mut^+^: GS115 *ΔHIS4*) and HSA-expressing (Mut^S^ ALB: GS115 *ΔHIS4 aox1*::*HSA*) cell cultures collected before, and at 3 and 24 hours after methanol induction (**Fig. 1a**). Ribo-seq uses a non-specific nuclease (*e.g.*, RNase A) to break down nucleic acids, including mRNA, that are not protected by ribosomes. To sequence ribosome protected mRNA fragments and reveal translational dynamics, ribosome derived RNA (rRNA) first needs to be depleted. We found that previous strategies to remove rRNA contamination in *K. phaffii* collected at log-phase growth in YPD media (Alva et al., 2021) were not sufficient for generating high quality Ribo-seq libraries where cells are collected at different growth stages and in different media. Our datasets agreed with previous Ribo-seq analyses (Ingolia et al., 2009) and revealed that a subset of rRNA represented the majority of rRNA contamination (**Supplementary Table 1**). Complementary oligos from pre-induction samples were used for probe-directed DSN treatment, reducing rRNA contamination from 88% to 10% in the pre-induction sample, 87% to 20% at 3 hours post-induction, and 93% to 62% at 24 hours post-induction.

**Fig. 1.**
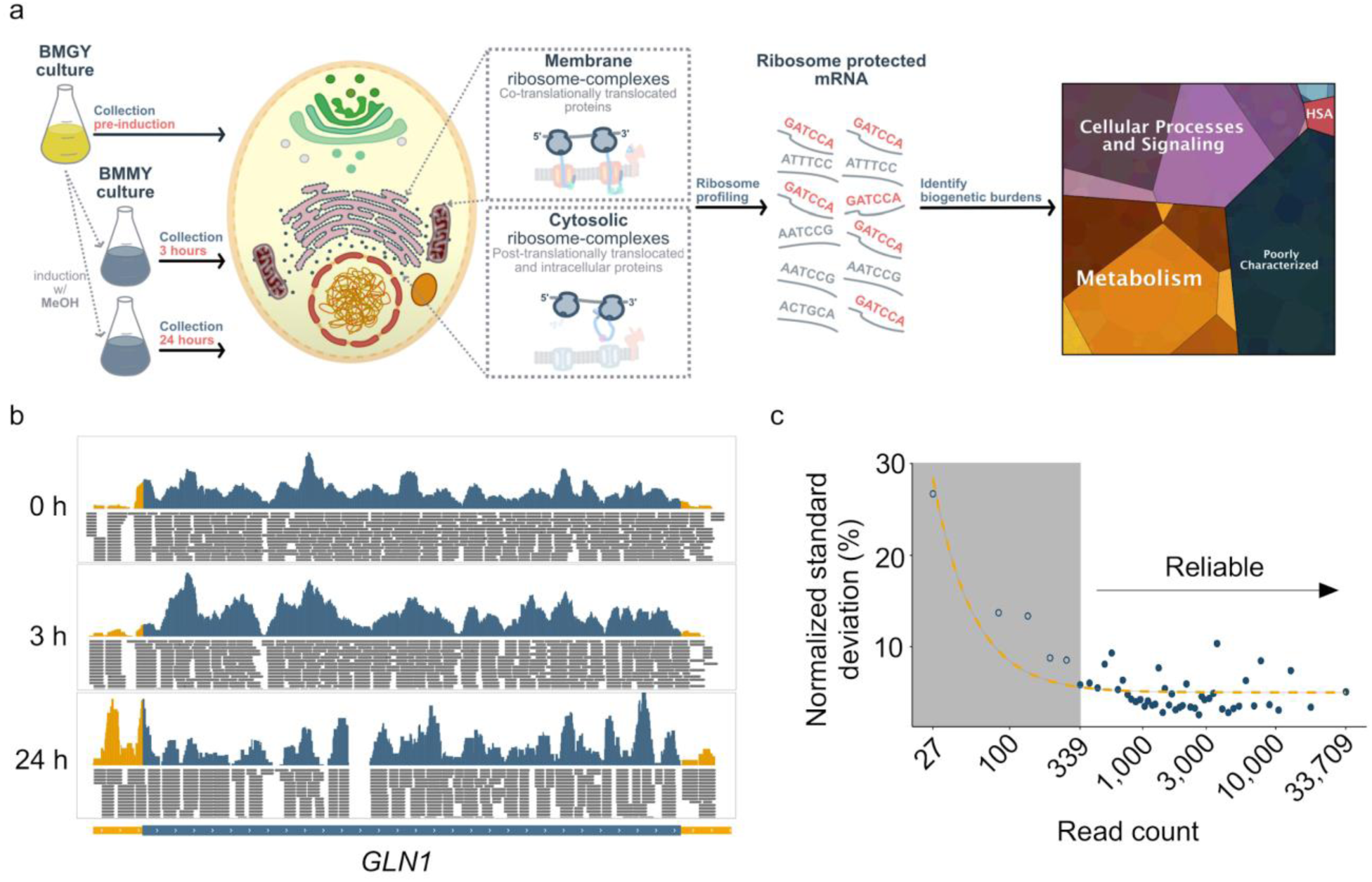
Ribo-seq analysis of protein expression in *K. phaffii*. a) Cultures were grown in buffered glycerol media (BMGY) to generate biomass. Cells were collected and transferred to buffered methanol media (BMMY) for heterologous induction. Samples were collected 3 and 24 hours post induction. Membrane-associated and cytosolic ribosomes were isolated, corresponding to co- and post-translationally translocated proteins, respectively. mRNA footprints were then isolated from ribosomes before Illumina sequencing. b) Ribosome abundance on transcripts under heterologous conditions for *GLN1*. The represented transcript reads are from GS115 Mut^S^ ALB cultures collected from the pre-induction sample cultured in BMGY media, 3 h and 24 h postinduction in BMMY media. The bottom blue band shows the ORF while the yellow band shows the UTRs. After 24 hours in the induction media a much higher proportion of reads map to the 5’UTR. Images are modified screen captures from Integrated Genome Viewer (MIT). c) Determining reads per gene thresholds. Biological replicates were used to determine read count thresholds when comparing genetic expression between data sets. Total read counts per gene were calculated by summing reads per gene for each replicate. Genes were binned according to this total read count value. Replicate read fractions were calculated by dividing read counts per gene by their bin value. Standard deviations of replicate read fractions were computed across each bin. Standard deviations were fit to replicate reads per gene using a generalized exponential decay model. Minimum read thresholds were calculated as the inflection point in this regressed curve. When reads per gene are fewer than this threshold, counting errors predominate inter-replicate variability. When reads per gene exceed this threshold, other sources of error predominate.

Before and three hours after induction, nearly all reads mapped to open reading frames (ORFs) and only 2% of reads mapped to untranslated regions (UTRs). Twenty-four hours after induction, however, we observed increased reads mapped outside of annotated ORFs as nearly 7% of reads mapped to UTRs. This was particularly true for genes like *GLN1* and *GCN4* that have previously been shown to have increased read density at 5’UTRs in response to stress (Ingolia et al., 2009) (**Fig. 1b**).

Our data revealed genome-wide coverage of expression, as up to 93% and 94% of *K. phaffii*’s 5,330 annotated protein-encoding genes were detected on average for GS115 Mut^+^ and GS115 Mut^S^ ALB, respectively (**Supplementary Fig. 1 and Supplementary Data 1 and 2**). Before making intra- and inter-sample comparisons of expression levels, we first sought to normalize reads (**Fig. 1c**). First, footprint sized fragments were used to generate computational masks for codons with a propensity to map to multiple locations of the genome. Next, reads per codon for the first 500 codons were normalized in all genes to account for positional counting biases in codons that were masked. Finally, we determined gene read count thresholds for comparing expressions between samples. To calculate these thresholds, we used biological replicates in the GS115 Mut^S^ ALB strain. In doing so, genes were binned according to the probabilistic distribution of the summed read counts per gene between each replicate. Binned genes’ read counts were normalized by the total read counts between both replicates. The standard deviation of each gene’s normalized reads with respect to their bin’s read count value was used to calculate read count thresholds necessary to reduce inter-replicate variability. These thresholds were used to predict read count thresholds for all samples as a function of their summed reads. This conservatively calculated read count thresholds between 52 reads to 573 reads, where samples with greater total reads had greater count thresholds. These criteria filtered approximately 1% of total nascent chains calculated per sample.

### 2.2 Translational landscape under heterologous conditions

We used Ribo-seq to survey nascent chain production in GS115 Mut^+^ and GS115 Mut^S^ ALB cultures collected before methanol induction and 3 and 24 hours after methanol induction. **Fig. 2a** shows the frequency distribution of log_2_ fold changes in gene expression relative to pre-induction levels. In both strains, gene expression undergoes significant changes at 24 hours compared to the 3-hour time point. While the expression levels of 125 and 209 genes remained unchanged (−0.025 < log_2_ fold change < 0.025) in GS115 Mut^+^ and GS115 Mut^S^ ALB three hours post-induction, these numbers decreased to 29 and 51 after 24 hours indicating a transcriptional response to methanol over the induction period.

**Fig. 2.**
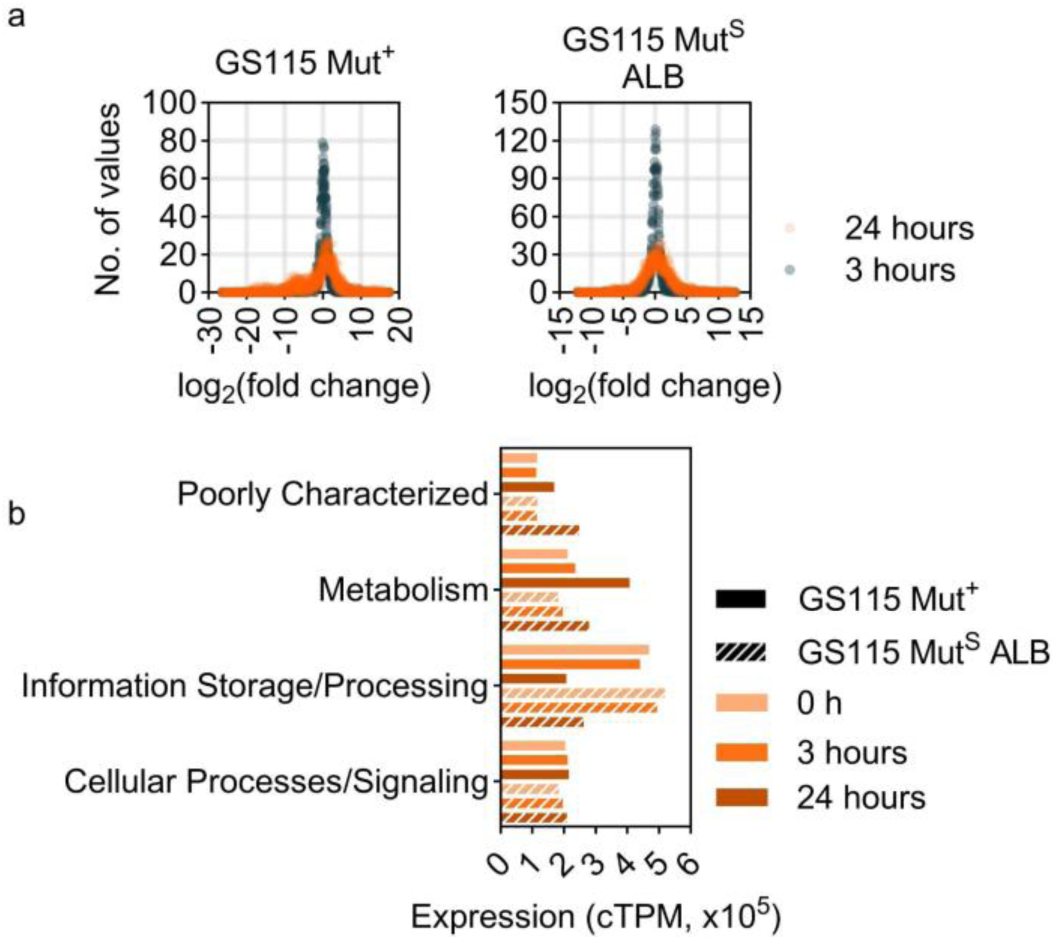
Ribo-seq analysis of gene expression pre- and post-induction. a) Gene expression fold change distribution 3 and 24 hours after methanol induction for GS115 Mut^+^ and GS115 Mut^S^ ALB strains. Most of the genes display constant levels of expression 3-hour post-induction, whereas gene expression levels dramatically change 24 hours after methanol induction for both strains. b) Total nascent chain production of genes belonging to cellular processes and signaling, metabolism, information storage and processing, and poorly characterized categories. Genes involved in information storage and processing experience reduced expression whereas genes in the cellular processes and signaling pathways remain their expression levels.

Next, we studied the total nascent chain production related to specific ontological categories including cellular processes and signaling, information storage and processing, and metabolism pathways as well as genes with functions that have yet to be characterized in each strain (**Fig. 2b**). The cumulative nascent chain production for genes involved in cell processes and signaling is relatively conserved; however, certain genes within this category, particularly those involved in signal transduction mechanisms, are notably upregulated (**Supplementary Fig. 2 and Supplementary Fig. 3**). Among these, genes related to methanol metabolism exhibit a substantial increase, such as the transcription factor *MIT1*, which shows a 148- and 4870-fold upregulation in the control and HSA-producing strains, respectively. This overexpression is linked to elevated expression of *AOX1* and *FDH1* genes, critical for methanol consumption and metabolism(Bachleitner et al., 2024).

In contrast, the total production of nascent chains related to information storage and processing decreased significantly by 56% in GS115 Mut^+^ and 49% in GS115 Mut^S^ ALB after 24 hours of methanol induction. This decrease was primarily driven by the reduced expression of genes involved in translation and ribosome biogenesis, which dropped by 73% in GS115 Mut^+^ and 57% in GS115 Mut^S^ ALB. Among these, several translation initiation factors (TIFs) are shown to be downregulated, in agreement with other studies (Rebnegger et al., 2014; Staudacher et al., 2022).

Uncharacterized genes showed increased expressions of 86% and 129% in GS115 Mut^+^ and GS115 Mut^S^ ALB strains after 24 hours of induction. The most differentially expressed uncharacterized proteins are those predicted to localize in the peroxisome, where energy is produced in the methanol utilization (MUT) pathway with a 4.2- and 3.4-fold increased expression in GS115 Mut^+^ and GS115 Mut^S^ ALB.

Total nascent chain production for metabolism-related genes increased by 81% and 45% in GS115 Mut^+^ and GS115 Mut^S^ ALB strains, respectively. Genes involved in the synthesis, transport, and catabolism of secondary metabolites, lipids, and carbohydrates are affected the most. Although amino acid transport and biosynthesis-related genes are not the most differentially expressed overall, we observe significant differential expression in specific genes, such as glutamine synthetase *GLN1*, which shows a 7.1-fold increase in GS115 Mut^+^ and 3.23-fold increase in GS115 Mut^S^ ALB, and proline oxidase *PUT1*, with a 670- and 6.1-fold increase in GS115 Mut^+^ and GS115 Mut^S^ ALB. The product of these genes import amino acids constituent of thiol-containing peptides involved in redox reactions, upregulated due to the presence of H_2_O_2_ (Farrugia and Balzan, 2012; Harding et al., 2003; Morotti et al., 2021).

### 2.3 Translational changes in methanol utilization and stress response pathways

Gene expression levels in the methanol utilization pathway shift under heterologous protein expression. While both strains showed increased expression of metabolic genes 24 hours after methanol induction, we observe that MUT pathway-related genes are expressed 3.9-fold more in GS115 Mut^+^ compared to GS115 Mut^S^ ALB (**Fig. 3**). This can be explained by higher methanol utilization rates in the control strain due to the presence of both *AOX1* and *AOX2*. The MUT pathway is divided into an assimilative branch, which produces biomass from formaldehyde, and a dissimilative branch, which generates CO_2_ from methanol and NADH to produce ATP(Cámara et al., 2017; Vogl et al., 2016; Yano et al., 2009a). As part of the assimilative branch, genes involved in the pentose phosphate pathway (PPP) also experience differential expression levels. Additionally, the reactive oxygen species (ROS) defense mechanisms are upregulated in the presence of methanol, as the oxidation of methanol to formaldehyde produces reactive hydrogen peroxide.

**Fig. 3.**
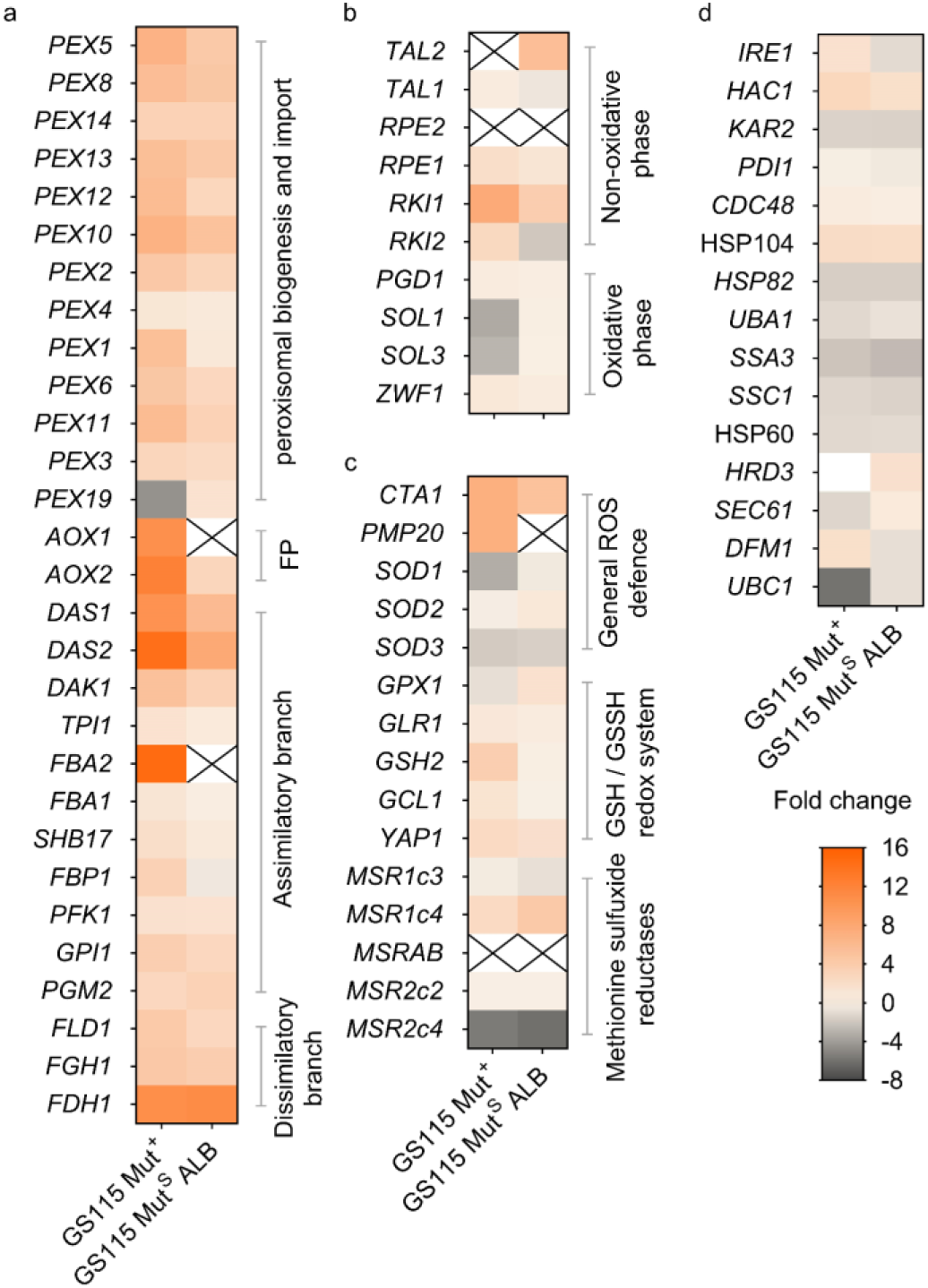
Regulation of methanol utilization and stress response pathways genes. Gene expression fold change for genes involved in a) Canonical methanol utilization pathway, b) Pentose phosphate pathway, c) Reactive oxygen species (ROS) pathway, and d) Unfolded protein response (UPR). Fold change is calculated as the log_2_ ratio of expressed genes 24 hours after methanol induction compared to the expression levels before induction.

We find a 58- and 19-fold increased expression of peroxisomal encoding genes 24 hours after growth in methanol media in GS115 Mut^+^ and GS115 Mut^S^ ALB, respectively (**Fig. 3a**). In the MUT pathway, AOX is produced exclusively in the presence of methanol and the absence of glucose, breaking down methanol into hydrogen peroxide and formaldehyde in the peroxisome. While there are two genes that encode for AOX in GS115 Mut^+^, the majority of AOX activity is expressed through *AOX1* as our datasets detect 32-fold greater expression from *AOX1* than *AOX2* after 24 hours of growth in the induction media similar to what is suggested in other studies (de Hoop et al., 1991; Zhang et al., 2009). Although GS115 Mut^S^ ALB has a deleted *AOX1*, we see a nearly 7-fold increase in the *AOX2* expression levels after 24 hours.

In the assimilative branch, formaldehyde reacts with xylulose 5-phosphate (Xu5P) produced in the PPP by dihydroxyacetone synthase (*DAS1* (ACN76559.1), *DAS2* (ACN76560.1), and possibly *TKL1*) (Krainer et al., 2012; Küberl et al., 2011). Our datasets show that translation of *DAS2* occurs more extensively than *DAS1* as it produces 15.8- and 3.7-fold more nascent chains compared to *DAS1* in GS115 Mut^+^ and GS115 Mut^S^ ALB strains after 24 hours. The cumulative expression of *DAS1* and *DAS2* is nearly 7 times higher in GS115 Mut^+^ compared to GS115 Mut^S^ ALB. The genes involved in the PPP related to biosynthesis of Xu5P are on average 46% and 39% upregulated in GS115 Mut^+^ and GS115 Mut^S^ ALB (**Fig. 3b**). Lower expression of genes to convert formaldehyde to other products in the GS115 Mut^S^ ALB strain reflects the slower methanol assimilation in this strain as a result of the knocked out *AOX1* which results in the slower production of formaldehyde.

Outside of the peroxisome, formaldehyde is dissimilated to formate by formaldehyde dehydrogenase (FLD) and to carbon dioxide by formate dehydrogenase (FDH) for energy production. We see a 16.8- and 10.1-fold increase in expression levels of FLD genes in GS115 Mut^+^ and GS115 Mut^S^ ALB, as well as a 1717.8- and 2179.7-fold increase for *FDH1*. Other studies have also shown that enzymes involved in the dissimilatory branch of the MUT pathway experience a drastic increase in response to heterologous production under methanol induction(Vanz et al., 2014, 2012). We see a 7.8- and 3.6-fold increase in the nascent chain production of mitochondrial malate dehydrogenase (*MDH1*) in GS115 Mut^+^ and GS115 Mut^S^ ALB 24 hour after methanol feeding likely due to the increased NADH supply from the methanol dissimilation branch and the heightened energy demand for growth, production, and cell maintenance, such as repair and recycling(Vanz et al., 2012).

Within the peroxisome, toxic hydrogen peroxide is decomposed into oxygen and water by catalase (*CTA1*) as the first step of the ROS defense. *CTA1* exhibits differential expression levels of 130.7-fold and 32.7-fold in GS115 Mut^+^ and GS115 Mut^S^ ALB (**Fig. 3c**). The lesser *CTA1* expression increase in the HSA-producing strain is attributable to its lack of a functional *AOX1* gene, resulting in a slower methanol assimilation rate and, consequently, reduced hydrogen peroxide production. As hydrogen peroxide causes oxidative stress, we also observed increased expression involved in oxidative stress responses for genes like *YAP1*, 5.3- and 3.5-fold increase in the GS115 Mut^+^ and GS115 Mut^S^ ALB strains, and *GSH2* with 12.6- and 1.14-fold increase in the GS115 Mut^+^ and GS115 Mut^S^ ALB strains.

Heterologous protein production in GS115 Mut^S^ ALB also changes the nascent chain production of genes involved in the UPR pathway (**Fig. 3d**). After induction, total nascent chain production of genes involved in UPR are decreased for both strains (18% and 30% reduction in the control and ALB-producing strain, respectively) suggesting that the HSA production likely has low impacts on UPR activation. The lower differential changes of many genes related to UPR in the ALB-producing strain including the *IRE1*, *HAC1*, and *KAR2*, the disulfide isomerase *PDI1,* the mitochondrial chaperones *HSP60* and *SSC1,* as well as cytosolic chaperones including *HSP104, CDC48, UBA1, SSA3* involved in disassembling and degrading the misfolded proteins indicate lower activation of the UPR/ERAD pathways in the ALB-producing strain compared to the control strain. This unexpected result has also previously been observed during methanol-induced production of insulin precursor in controlled fed-batch cultures(Vanz et al., 2014), (Roth et al., 2018). It has been reported that cells cultured in glycerol media experience higher levels of UPR activation. This suggests that the elevated UPR activity might help alleviate the burden associated with the strong induction of the recombinant product(Kastberg et al., 2022).

### 2.4 Heterologous expression and host protein biogenesis demands

We next compared host protein synthesis in GS115 Mut^+^ and GS115 Mut^S^ ALB cultures. Prior to induction, protein synthesis rates per gene are highly conserved between GS115 Mut^+^ and GS115 Mut^S^ ALB as they have a Pearson’s correlation of 0.97 (**Fig. 4a, Supplementary Fig. 4**). However, expression diverges significantly over time between the two strains as they show Pearson’s correlations of 0.94 and 0.32 after 3 and 24 hours of methanol induction.

**Fig. 4.**
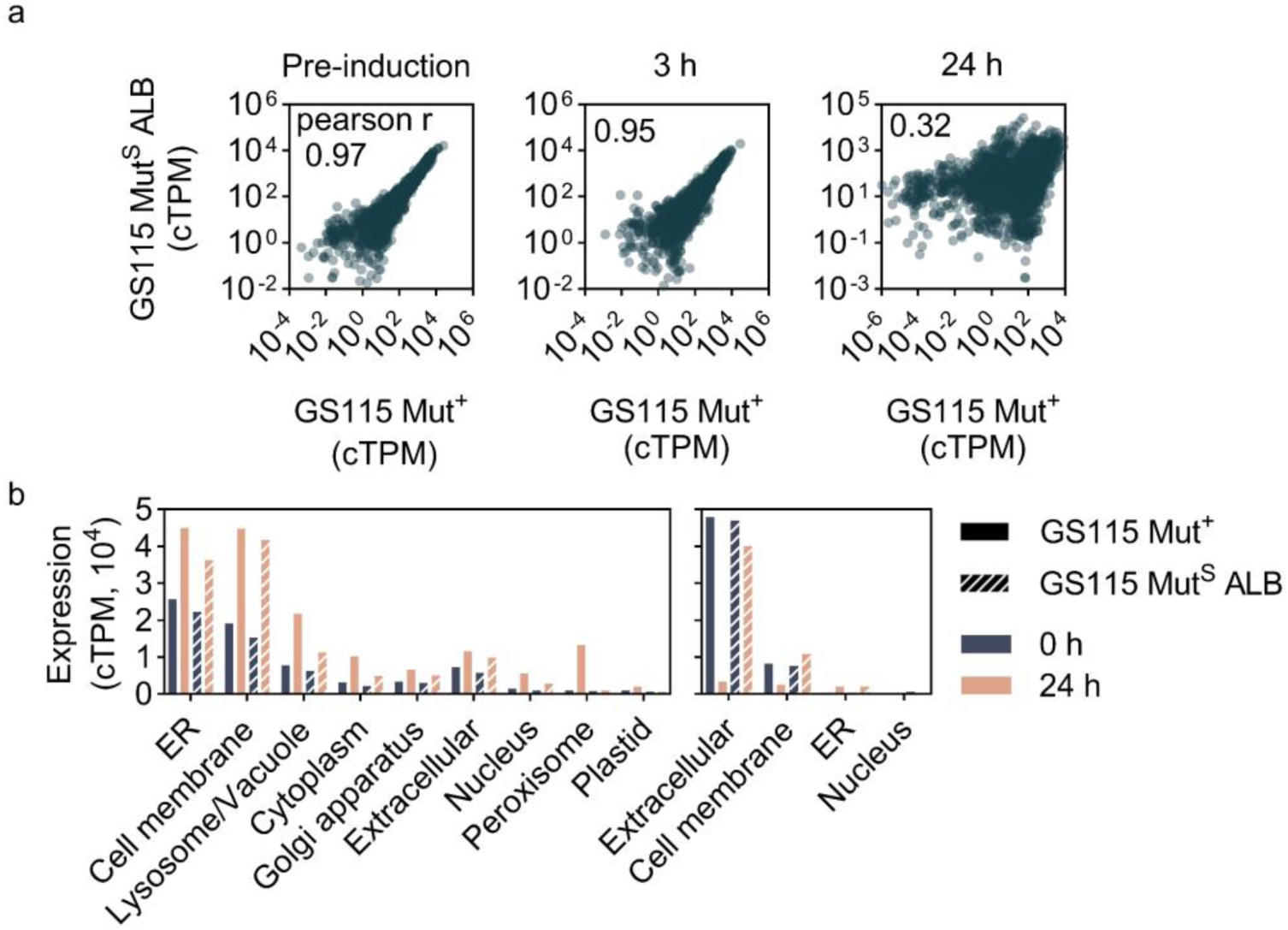
Divergence of translational landscape after heterologous expression. a) Prior to heterologous induction, nascent proteins produced per gene for GS115 Mut^+^ and GS115 Mut^S^ ALB are highly correlated. After heterologous induction with methanol media, expression diverges between the two strains. b) Total nascent chain production of proteins translocating into the ER co-(left) and post-translationally (right) belonging to various subcategories. In the albumin-producing strain, proteins localized to the cell membrane through co- and post-translational mechanisms exhibited the highest increase in expression 24 hours after induction.

Host cell proteins can translocate into the ER using either co- or post-translational pathways, while heterologous proteins typically use co-translational pathways. In *K. phaffii*, we estimate 56 protein products to enter the ER post-translationally and 931 protein products to enter the ER co-translationally. Before induction, about 12% of nascent chains in each strain were predicted to enter the secretory pathway, with a similar distribution between co- and post-translational entry into the ER (**Supplementary Fig. 5**). This distribution aligns with findings from our previous study on *K. phaffii(Alva et al., 2021)*. After 24 hours of methanol induction, this increased to around 17% for both strains. However, the ratio of nascent chains entering the ER co- and post-translationally greatly diverges between strains. Post-translational translocation decreases significantly in GS115 Mut^+^ 24 hours after induction, with nearly an 85% reduction, while it remains unchanged in GS115 Mut^S^ ALB. The total production of post-translationally translocated nascent chains by GS115 Mut^S^ ALB was 6.6 times higher compared to the control strain, with the majority of these nascent chains being extracellular (27 genes) and membrane proteins (16 genes) (**Fig. 4b)**. Co-translational translocation, however, increases in both strains, with a 2.3-fold increase in GS115 Mut^+^ and a 2-fold increase in GS115 Mut^S^ ALB. While co-translationally translocated proteins localized to the cell membrane experience the most increase for the HSA-producing strain, peroxisomal proteins in the control strain show the greatest increase, likely due to faster methanol consumption.

Both co- and post-translationally translocated genes experience different expression levels between the two strains (**Fig. 5** and **Supplementary Fig. 6**). We investigated which host cell proteins might limit heterologous protein entry into the ER by sequestering Sec-translocons during different stages of expression in GS115 Mut^S^ ALB. To focus on proteins that could restrict bioproduction, we excluded ER-resident proteins from our “hit list” as their deletion could impair folding and secretion of heterologous proteins. After methanol induction, the most highly differentially expressed proteins between the two strains that are co-translationally translocated are *ATO2*, *YDR134C*, *GAL2*, and *BGL2. ATO2* is a putative transmembrane protein involved in export of ammonia and it is the paralog of *ADY2* in *S. cerevisiae*; this gene was not targeted in our strain engineering due to its fitness effects (Tafrishi et al., 2024). *YDR134C, GAL2,* and BGL2 were all selected for knockout. *YDR134C* is a secreted protein involved in cell wall maintenance that is homologous to *S. cerevisiae*’s paralog of *CCW12*. *GAL2* is a membrane protein involved in carbohydrate import and acetate transport. *BGL2* is endo-beta-1,3-glucanase and is a major protein of the cell wall involved in cell wall maintenance.

**Fig. 5.**
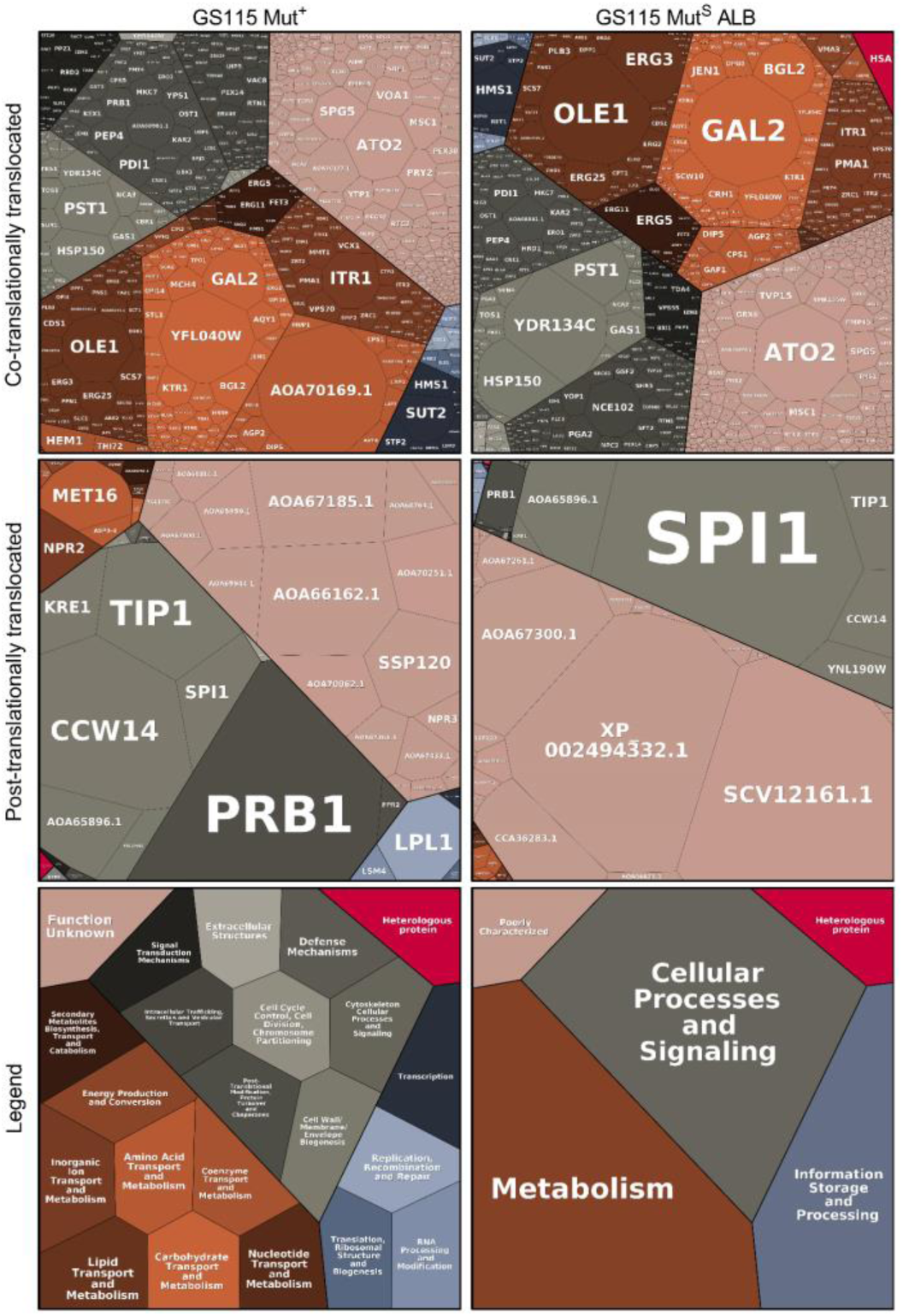
Co and post-translational flux through the ER for GS115 Mut^+^ and GS115 Mut^S^ ALB strains 24 hours after induction. Non-mitochondrial proteins are predicted to enter the secretory pathway co-translationally if they have greater than log_2_ membrane enrichment in YPD studies. Gene products are grouped by ontological function using COG scores predicted by EggNOG v5.0. Cell sizes are calculated using cTPM scores and represent relative quantities of nascent chains produced per gene. Tessellation plots are made using www.bionic-vis.biologie.uni-greifswald.de(Bernhardt et al., 2009; Liebermeister et al., 2014; Otto et al., 2010).

For nascent chains that enter the ER post-translationally, the most highly expressed proteins are secreted and remain conserved before and after induction. The most highly differentially, post-translationally translocated proteins are *SPI1*, *XP_002494332.1*, *SCV12161.1,* and *AOA65896.1*. These proteins are relatively small cell wall constituents, and they are predicted to localize extracellularly. *SPI1* has been identified as an essential gene in *K. phaffii* by our functional genomic screening and therefore is removed from the target list(Tafrishi et al., 2024). Our final curated list includes *YDR134C*, *GAL2*, *BGL2, XP_002494332.1*, *SCV12161.1,* and *AOA65896.1,* all of which are non-essential genes that enter the early secretory pathway and are highly differentially expressed between the control and heterologous protein producing strain. We hypothesize that by knocking out these genes we may be able to rationally engineer strain with improved protein secretion.

### 2.5 Ribo-seq-enabled rational engineering of *K. phaffii* to improve protein secretion

Knocking out highly expressed, non-essential genes sequestering the most resources related to protein secretion under heterologous conditions is a rational approach for genetic modification. To test this approach, we created single and multiple gene knockouts in GS115 Mut^S^ ALB cells in *GAL2*, *YDR134C*, *BGL2*, *SCV12161.1*, and *AOA65896.1* genes. We were unsuccessful in knocking out *XP_002494332.1* even with multiple sgRNAs and therefore this gene was removed from further analysis.

To measure HSA secretion, engineered cells were first cultivated in glycerol media for biomass accumulation, then transferred to buffered methanol media including 0.5% methanol, with daily addition of 0.5% methanol. After five days, the supernatant was collected, concentrated for better visualization of bands on SGS-PAGE gels, and the HSA band volume was measured. (**Fig. 6a**). For accurate secretion measurements, first the band volume of HSA on each lane was normalized to the OD_600_ of the sample in order to be able to differentiate between the impact of gene knockouts on the fitness of the cell and HSA secretion levels. HSA band volume was also normalized to the band volume of blue fluorescent protein (BFP) added equally to each well to standardize the band volumes enabling a more accurate comparison between lanes. The relative normalized band volume was calculated as the ratio of normalized band volume of each sample to the same of non-engineered cells (GS115 Mut^S^ ALB) (**Fig. 6b and c**).

**Fig. 6.**
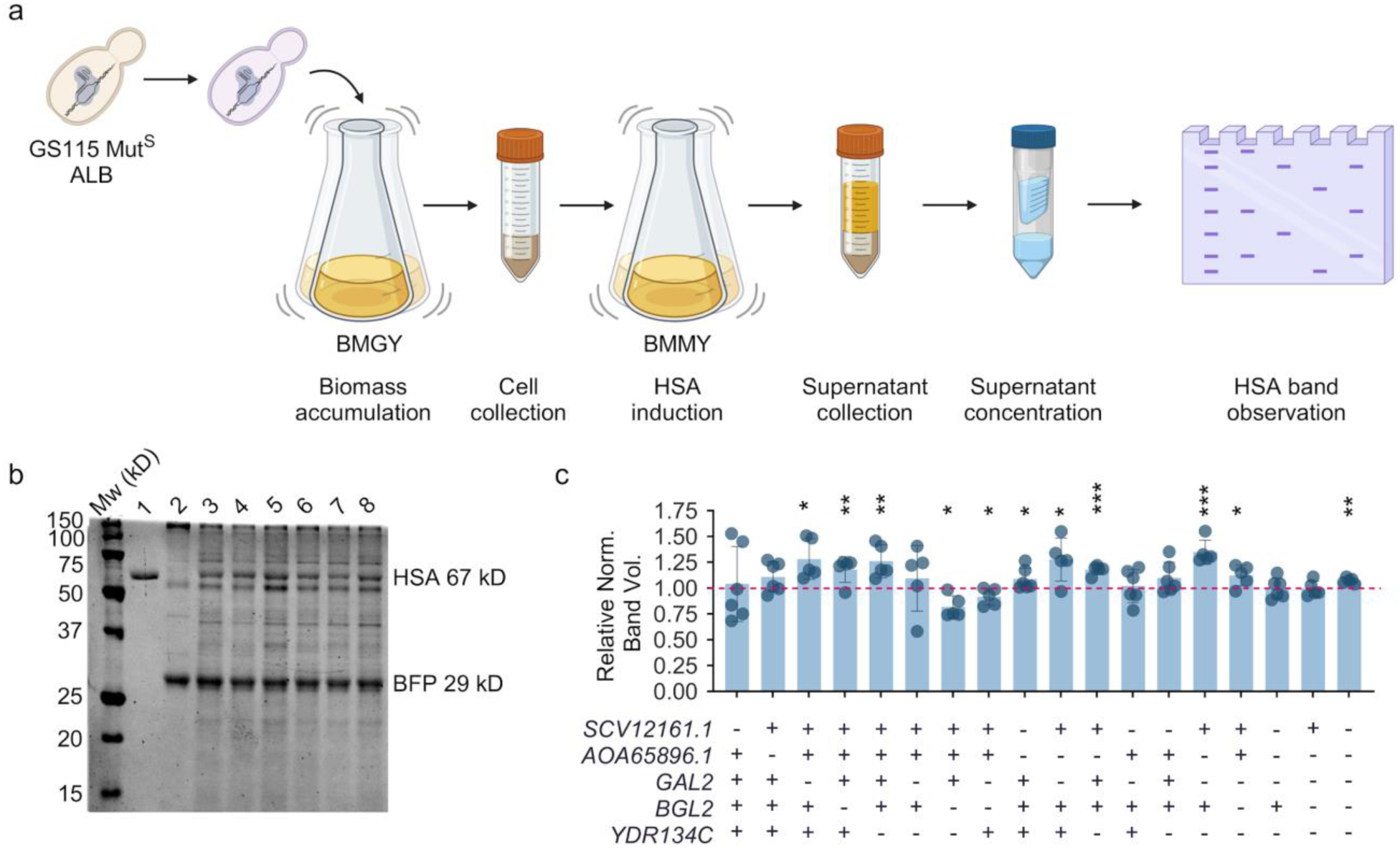
Ribo-seq guided strain engineering. a) Experimental workflow of rational engineering of *K. phaffii* cells enabled by Ribo-seq. Five genes with high expression levels sequestering the early secretory pathway were identified by Ribo-seq. Subsequently single and multiple knockout strains were produced using the CRISPR-Cas9 system in the GS115 Mut^S^ ALB strain. Strains were grown in buffered glycerol media (BGMY) overnight and then transferred to buffered methanol media (BMMY) for HSA induction for five days. The supernatant was extracted, concentrated using filters with a 50 kDa cutoff, and mixed with Blue Fluorescent Protein (BFP) as the standard allowing for a more accurate HSA qualification between lanes. HSA band volume was normalized and measured on SDS-PAGE gels to compare heterologous protein secretion. b) Secreted proteins into the media from GS115 Mut^S^ ALB cells with single knockouts visualized on SDS-PAGE gels. Lanes from left to right: protein standard, pure recombinant HSA (1), GS115 Mut^+^ strain (2), GS115 Mut^S^ ALB (3), GS115 Mut^S^ ALB *gal2* (4), GS115 Mut^S^ ALB *aoa65896.1* (5), GS115 Mut^S^ ALB *bgl2* (6), GS115 Mut^S^ ALB *scv12161.1* (7), GS115 Mut^S^ ALB *ydr134c* (8). c) Relative normalized HSA band volume in strains with either single or multiple gene knockouts. Bars represent the ratio of secreted normalized HSA from each knocked out strain compared to the base GS115 Mut^S^ ALB strain. HSA band volume was normalized to the OD_600_ of each sample and the band volume of blue fluorescent protein (BFP) added equally to each lane. Experiments were done in six biological replicates and the bars show the mean of the replicates. * p < 0.05, ** p < 0.005, and *** p < 0.005; one-tailed paired t-test).

We first investigated the influence of single knockouts on HSA secretion. It was observed that a single knockout in co-translationally translocated *GAL2*, *BGL2*, and *YDR134C* genes was able to increase HSA secretion 28, 18, and 26%, respectively, compared to the control strain. Whereas single knockouts in post-translationally translocated *SCV12161.1* and *AOA65896.1* genes did not statistically change HSA secretion. This might be attributed to the fact that HSA is being translocated into the ER co-translationally and alleviating the load on these chaperones is able to increase HSA secretion.

We next sought out the influence of double gene knockouts on HSA secretion. The addition of *BGL2* knockout to strains with either *YDR134C* or *GAL2* single knockout was associated with statistically significant lower levels of HSA secretion (0.82- and 0.92-fold secretion in GS115 Mut^S^ ALB *ydr134c bgl2* and GS115 Mut^S^ ALB *gal2 bgl2* double-knockouts, respectively, compared to the non-engineered strain). Furthermore, HSA secretion did not change with the addition of *AOA65896.1* knockout to the GS115 Mut^S^ ALB *gal2* strain, and it resulted in lower secretion levels in the GS115 Mut^S^ ALB *ydr134c* strain. The highest increase in HSA secretion was observed with a triple-knockout in *GAL2, AOA65896.1* and *YDR134C* genes which led to a 35% improvement of HSA secretion compared to the control strain. Other studied combinations or additional knockouts were not able to improve secretion further.

## 3. Discussion

Understanding how production strains operate under heterologous conditions can provide insights into optimizing bioproduction (Kroukamp et al., 2017). To this end, we compared host protein synthesis between a control (GS115 Mut^+^) and a recombinant protein producing (GS115 Mut^S^ ALB) *K. phaffii* strain under heterologous conditions. We used Ribo-seq to understand how recombinant protein production affects early secretory trafficking of host cell proteins under heterologous conditions. Highly expressed host cell proteins that enter the early secretory pathway sequester biogenesis machinery that are limited in number and processivity which may limit heterologous secretion. Host cell proteins may enter the early secretory pathway co- or post-translationally. We previously used protein sequence features as well as Ribo-seq reads from cytosolic and membrane bound ribosomes in GS115 Mut^+^ cultured in YPD to predict the trafficking pathways of these proteins (Alva et al., 2021). The assumption that proteins translocate similarly under heterologous conditions relies on two previous observations: *K. phaffii*’s secretome does not change with different carbon substrates (Burgard et al., 2020) and that proteins’ ER translocation routes depend on their sequence features and size (Alva et al., 2021; Chartron et al., 2016).

In comparing GS115 Mut^+^ and GS115 Mut^S^ ALB, the percentage of nascent chains predicted to enter the ER increased after 24 hours of methanol induction. However, a significantly greater number of cell wall and membrane nascent chains entered the ER for GS115 Mut^S^ ALB. The molecular organization of the cell wall is dynamic. The mechanical strength of the cell wall is largely due to the inner layer consisting of β 1,3-glucan and chitin (Klis et al., 2002). The outer layer of the cell wall consists of glycosylated mannoproteins covalently linked to the β 1,3-glucanchitin network directly or disulfide bound to other cell wall proteins. Cell wall mannoproteins affect stability and resistance to stress (Mrsa et al., 1999; Shimoi et al., 1998; van der Vaart et al., 1995). Because the differences in extracellular and membrane proteins between strains do not appear to result from oxidative stress-induced expression (Thorpe et al., 2004), the reorganization of the cell wall in the GS115 *Mut^S^* ALB strain is likely due to the stress caused by heterologous protein secretion. Therefore, the most highly expressed cell wall and membrane proteins entering the ER at different stages of heterologous expression offer novel insights for improving secretion. The effect of cell wall proteins on recombinant protein secretion is not well studied. A recent study showed the elevated expression levels of green fluorescent protein (GFP) and human epidermal growth factor (hegf) as well as looser cell surface and larger cell size due to inactivation of two genes involved in cell wall β-glucan biosynthesis including *FSK1* and *FSK2 (Cheng et al., 2024)*. Another study found the disruption of the cell wall mannoprotein *CWP2* increased cell wall permeability and cellobiohydrolase heterologous secretion (Li et al., 2022, 2020; Zhang et al., 2008). Another study found the inactivation of genes involved in cell wall biosynthesis including *DFG5*, *YPK1*, *FYV5*, *CCW12* and *KRE1* increased cellulolytic enzyme β-glucosidase secretion and surface display in *S. cerevisiae* (Chen et al., 2023).

While the diversity and number of post-translationally translocated nascent chains do not appreciably change after induction, their expression levels are amongst the highest observed. Of this group, three small, hypothetical cell wall proteins including *XP_002494332.1*, *SCV12161.1,* and *AOA65896.1* are amongst the most highly differentially expressed. The upregulation of *AOA65896.1* and *XP_002494332.1* compared to the control strain has also been observed in the production of lysozyme using chemostat cultures(Hesketh et al., 2013). For co-translationally translocated gene products, *GAL2*, *YDR134C*, and *BGL2* are amongst the most highly differentially expressed between strains. As galactose is preferentially incorporated into cell wall glucan over glucose (Coen et al., 1994), we speculate that *GAL2* is upregulated to increase the overall expression of cell wall mannoproteins. Since the carbon source used in these experiments does not contain galactose, *GAL2* is a good candidate for strain engineering under heterologous conditions. *YDR134C* and *BGL2* are both major cell wall proteins. *YDR134C* is a heavily glycosylated, nonenzymatic GPI-anchored protein involved in the cell wall organization. *BGL2* is involved in cell wall maintenance and incorporating newly synthesized mannoprotein molecules into the cell wall. We used the CRISPR-Cas9 system to functionally knockout the identified genes. Single knockouts of Gal2, Ydr134c, and Bgl2 enhanced HSA secretion, while functionally inactivating the post-translationally translocated genes had no significant effect. Double knockouts involving *BGL2* reduced secretion, highlighting potential functional dependencies. The highest improvement (35%) was achieved with a triple knockout of *GAL2*, *YDR134C*, and *AOA65896.1*, demonstrating the potential of cell wall engineering for optimizing protein production in *K. phaffii*.

While RNA-seq has been used to study oxidative stress responses proceeding methanol metabolism (Yano et al., 2009b), Ribo-seq is a more sensitive and appropriate tool for quantifying protein levels as oxidative stress increases the frequency of post-transcriptional modifications (Blevins et al., 2019; Gerashchenko et al., 2012; Yano et al., 2009b). For instance, RNA-seq finds *DAS1* and *DAS2* equally expressed after methanol induction (Krainer et al., 2012) while Ribo-seq shows *DAS2* to be more highly expressed than *DAS1*. Ribo-seq also reveals translational dynamics that indicate methanol induced oxidative stress responses. At many loci, we observe translation initiation events upstream ORFs at 5’UTRs after methanol induction similar to other studies of H_2_O_2_ treated yeast cultures (Gerashchenko et al., 2012). As well, our analyses are congruent with previously observed reductions in protein synthesis rates consequential to oxidative stress (Shenton et al., 2006) as we find decreased expression of genes encoding ribosomal proteins (e.g., *RPS23B* and *RPL3*) and increased expression of genes encoding RNA-binding proteins thought to stabilize slowly translating transcripts from degradation (e.g., *NRD1*, *NAB3*, and *PAB1*) (Vogel et al., 2011). Together, we find GS115 Mut^S^ ALB less affected by methanol induced oxidative stresses than GS115 Mut^+^, likely due to lesser AOX expression and subsequently lesser H_2_O_2_ generation. Additionally, we found that GS115 Mut^S^ ALB shows lower overall expression levels for genes involved in the UPR and ERAD compared to GS115 Mut^+^, similar to another study(Kastberg et al., 2022; Vanz et al., 2014).

This study provides valuable insights into how *K. phaffii* responds to heterologous protein production at the translational level, particularly in relation to early secretory trafficking and cell wall dynamics. Using Ribo-Seq, we identified key host cell proteins that sequester biogenesis machinery, potentially limiting recombinant protein secretion. Our findings highlight the significant role of cell wall and membrane proteins in shaping the secretion capacity of *K. phaffii.* Additionally, we observed lower expression of genes involved in the UPR and ERAD pathways, suggesting that the strain’s protein folding and degradation mechanisms are less affected under heterologous conditions. These findings advance our understanding of *K. phaffii*’s cellular adaptation to recombinant protein production and provide a foundation for strain engineering strategies aimed at optimizing industrial protein secretion.

## 4. Conclusion

Heterologous protein production is a complex process that relies on the limited resources of the translational and secretory machinery. Through next-generation sequencing and ribosome profiling, we have gained valuable insights into the metabolic and secretory demands of the methylotrophic yeast *Komagataella phaffii* under heterologous conditions. Our findings revealed host protein flux changes through the endoplasmic reticulum in response to heterologous protein production. We also found that the variety and levels of host proteins entering the secretory pathway are unique to different stages of heterologous expression. Application of this tool allowed us to find highly expressed, non-essential gene targets that consume resources in the early secretory pathway during methanol induction. Employing the CRISPR-Cas9 system, we performed single or combined gene knockouts to investigate the influence of these genes on rationally improving recombinant protein secretion in *K. phaffii*. Our investigation resulted in the identification of a triple knockout leading to a 35% improvement of human serum albumin (HSA) secretion from this industrially relevant yeast. These findings offer valuable insights for process optimization and strain engineering in industrial applications

## 5. Materials and Methods

### 5.1 Strains and culture conditions

All strains used in this work are presented in **Supplementary Table 2**. All experiments were performed using GS115 Mut^+^ and GS115 Mut^S^ ALB (Pichia expression kit, Life Technologies, 2014). For Ribo-seq experiments, 200 mL liquid cultures of BMGY (1 % yeast extract, 2 % peptone, 100 mm potassium phosphate pH 6.0, 1.34 % YNB, and 1 % glycerol) were grown to an OD_600_ nm of 5 at 30 °C with shaking in baffled 2 L flasks. Of this culture, 100 mL were harvested by vacuum filtration through a 0.8 µm filter. Immediately after filtering, cells were scraped off the filter using a chilled scoopula and submerged in a 50 mL conical tube containing liquid nitrogen. The remaining liquid cultures were split into two 50 ml conical tubes and were pelleted via centrifugation. Supernatant was removed from each 50 ml conical tube. The cell pellet of one 50 ml conical tube was gently resuspended with 40 mL BMMY without methanol (1 % yeast extract, 2 % peptone, 1.34 % YNB, and 100 mm potassium phosphate pH 6.0). Resuspended culture was used to resuspend the cell pellet in the second 50 ml conical tube. Resuspended cultures were equally divided into two 280 mL cultures of BMMY without methanol in 2 L baffled flasks for a final volume of 300 mL for each sample. Methanol was added at 0.5 % to each of the baffled flasks for *AOX1* induction. Flasks were allowed to shake at 30 °C and were collected in the manner described above three and twenty-four hours after methanol induction. Lysis buffers (50 mM MOPS, 25 mM KOH, 100 mM KOAc, 2 mM MgOAc, 1 mM DTT, and 1 % Triton X-100) for each sample were frozen by adding 2 mL dropwise to a 50 mL conical tube containing liquid nitrogen. For each sample, frozen cells were mixed with 2 mL frozen lysis buffer. Cell fractions were pulverized for 2 min in a 50 mL ball mill chamber with a single 2 cm steel ball (Retsch) and collected in 50 mL conical tubes. After thawing, lysates were centrifuged at 18,000 g for 10 min. Supernatants were transferred to 1.5 mL conical tube and were further clarified by centrifugation at 23,000 g for 20 min.

For yeast transformation experiments, cells were grown in 100 ml YPD at 30 °C and 225 RPM and were transformed with plasmids expressing sgRNAs. Transformants were grown in 2 ml of YPD supplemented with 800 μg/ml of G418 in 14 ml polypropylene tubes. For HSA secretion quantification, cells with the knockout of interest were grown in 200 ml of BMGY supplemented with 1% glycerol in 1 L shake flasks for biomass accumulation until OD_600_ = ∼ 6. Next, cells were transferred to 20 ml of BMMY with 0.5% methanol for HSA induction in 100 ml shake flasks. Cells were grown in BMMY media for five days with daily supplementation of 0.5% methanol (v:v).

### 5.2 Ribo-seq

Lysed samples were digested using 40 U of Ambion RNase A for one hour at room temperature. Digested samples were layered on a 10 % to 50 % sucrose gradient prepared in 50 mM Tris pH 7.5, 200 mM NaCl, and 2 mM MgOAc case using a Gradient Master (Biocomp). Gradients were centrifuged at 39,000 RPM for 2.5 h in a TH-641 rotor (Thermo). After centrifugation, gradients were fractionated using a Piston Gradient Fractionator (Biocomp) and monosome peaks were retained. Total RNA was extracted using a standard phenol-chloroform method and alcohol precipitated. Ribosome protected footprints 18 nt to 34 nt were resolved and excised using 15 % polyacrylamide TBE-urea gel. RNA was collected from excised gel fragments using RNA gel extraction buffer (300 mM NaOAc, 1 mM EDTA, and 0.25 % SDS), precipitated, and resuspended in water containing 20 U mL^−1^ SUPERase·In.

Purified fragments were then dephosphorylated by incubating 2 µL 1 M RNA sample with 2 µL RNase free water, 0.5 µL SUPERase·In RNase Inhibitor, 0.5 µL T4 Polynucleotide Reaction Buffer (PNK), and 0.5 µL T4 Polynucleotide Kinase at 37 °C for 1 h. Dephosphorylated samples were linker ligated with adapter sequences by incubating with 3.5 µL 50 % PEG-8000, 0.5 µL 10X T4 RNA Ligase Reaction Buffer, 0.5 µL 10 µM adenylated linkers and 0.5 µL T4 Rnl2(tr)k277Q at 30 °C for 4 h. Linker-ligated samples were concentrated via isopropanol precipitation and resolved using 15 % TBE-urea polyacrylamide gel. Imaged samples were diluted and pooled to equivalent concentrations by their relative pixel intensities calculated from BioRad imaging software after overnight extraction in RNA gel extraction buffer.

Ligated and purified samples were rRNA depleted using streptavidin-coated magnetic beads from the Ribo-Zero rRNA Removal Kit as recommended by the manufacturer. Depleted samples were precipitated, resolved using 15 % TBE-urea polyacrylamide gel, and extracted as previously described.

RNA was reverse transcribed by adding 2 µL reverse transcription primer to 10 µL sample and incubating at 65 °C for 5 min to denature. Denatured sample was then incubated with 4 µL 5X First Strand Buffer, 1 µL 10 mM dNTPs, 1 µL 10 mM DTT, 1 µL 20 U µL −1 SUPERase·In and 1 µL 200 U µL^−1^ SuperScript II Reverse Transcriptase at 50 °C for 30 min using thermal block. After incubation, the sample was hydrolyzed by adding 2.2 µL 1 M NaOH and then incubated at 70 °C for 20 min using a thermal block. 28 µL RNAse free water was added to reverse transcription mixture (∼50 µL total) and concentrated using the Oligo Clean and Concentrator Kit. Concentrated RNA was then purified of reverse transcription primers using 12 % TBE-urea polyacrylamide gel. RNA from gel slices was extracted using the method previously described. Extracted precipitants were resuspended in 11 µL 1:1000 SUPERase·In.

Single stranded cDNA samples were circularized by incubating 11 µL sample in 2 µL CircLigase II 10x Reaction Buffer, 1 µL 50 mM MnCl_2_, 1 µL ATP, 4 µL 5 M Betaine, and 1 µL 100 U µL^−1^ CircLigase II ssDNA Ligase at 60 °C for 3 h on thermal block. The circularization process was inactivated by incubating the sample at 80 °C for 10 min on a thermal block.

Circularized samples were rRNA depleted, again, using probe-directed degradation via double stranded nuclease (DSN) (Archer et al., 2015; Chung et al., 2015). Depletion probes were designed using rRNA aligned Ribo-seq reads collected from GS115 Mut^S^ ALB cultured in BMGY before methanol induction. Circularized samples (10 µL) were incubated with 4 µL 4x hybridization buffer, 1 µL 4x depletion probes at 200 µM, and 1 µL water. Mixture was denatured at 98 °C for 2 min and allowed to slowly anneal at 65 °C for 5 h. Double stranded rRNA fragments were enzymatically degraded by adding 2 µL 10x DSN master buffer, 1 µL DSN storage buffer, and 1 µL DSN before incubation at 65 °C for 25 min. Reaction was stopped by adding 20 µL 10 mM EDTA to DSN depleted sample mix. Samples were then purified using AMPure XP beads. After DSN treatment, samples were digested using Exonuclease I to degrade linearized DSN degraded DNA fragments as these may contain regions complementary to PCR amplification primers. Samples were again purified using AMPure XP beads.

Circularized samples were PCR amplified for 16 cycles using a 50 µL reaction mixture (10 µL Q5 Reaction Buffer, 1 µL 10 mM dNTPs, 2.5 µL 10 µM forward primer, 4 µL circularized DNA sample, 0.5 µL Q5 High Fidelity DNA Polymerase and 29.5 µL RNAse free water) divided into 5 x 10 µL aliquots. Amplified sample was resolved using 10 % non-denaturing TBE polyacrylamide gel and extracted using previously described method. Libraries were quantified using Qubit 2.0 Fluorometer and sequenced using Illumina NextSeq.

### 5.3 Mapping of ribosome-protected reads to codons

Sequenced reads were trimmed and demultiplexed in an error-tolerant way using Cutadapt 2.3 (Martin, 2011) (**Supplementary Table 3**). Reads were computationally rRNA subtracted by aligning them to *Komagataella pastoris* GS115 genomic rRNA using HISAT2 (Jain et al., 2015; Kim et al., 2015). Subtracted reads were mapped to the genome for *Komagataella pastoris* GS115 (Love et al., 2016) using HISAT2. Sequence alignment map (SAM) files were converted to sorted and indexed binary alignment map (BAM) files using Samtools and only included reads of high mapping quality (Danecek et al., 2021; Li et al., 2009). Genomic alignments were loaded into R using the GenomicAlignments package from Bioconductor (Lawrence et al., 2013). Genomic alignment ranges were converted to their 3’ end positions before determining p-site offsets. P-site offsets were determined using the existing genome annotations (Alva et al., 2021) and the RiboProfiling package in Bioconductor (Popa et al., 2016). Genomic alignment objects were used with p-site offsets to generate reads per codon per gene (RPCPG) data tables.

### 5.4 Masking reads of ambiguously mapped codons

Codon masks were created by first parsing the coding sequence annotation file associated with the reference genome into a FASTA file simulating every possible 28 NT combination (approximate length of a ribosome protected mRNA fragment). This FASTA file was then aligned to the reference genome twice, once to only include reads with mapping quality greater than or equal to 60 (unambiguously assigned), and another to include all reads (ambiguously assigned). Both alignment files were used to generate RPCPG data tables using methods previously discussed. The unambiguously assigned reads were subtracted from ambiguously assigned reads and codons with a nonzero difference were included in the mask. The first and last five codons in genes’ open reading frames were masked to correct for variable read quality at the beginning and ending of transcripts inherent to Ribo-seq (Mohammad et al., 2019).

### 5.5 Normalization and differential expression analysis

Read counts were normalized at the codon level using a metagene correction strategy previously discussed (Alva et al., 2021) with some modification. Reads for the first 500 codon positions at the 5’end of all transcripts was scaled by their respective codon-specific normalized metagene values. Normalized metagene values were calculated for all codons in all ORFs and applied in the following manner: positions 1 to 100 were normalized with a rolling mean with a window of 10 codons and positions 100 to 500 were normalized with a rolling mean with a window of 100 codons. Scaled reads per gene were calculated as the sum of a gene’s scaled codon reads (codon positions less than or equal to 500) and unscaled codon reads (codon position greater than 500).

Gene read count thresholds were calculated using an adapted method (Ingolia et al., 2009). First, we summed the scaled reads per gene for each gene between biological replicates. Each gene was grouped into 1 of 50 quantiles using the probabilistic distribution of the summed scaled read counts between replicates. In calculating the read count threshold for one replicate, the replicate’s scaled reads per gene were normalized by the summed read count for their respective bin. The standard deviation of normalized fractions across each bin were plotted against the summed read value for each bin. Read count thresholds were calculated as the knee-point in the exponential regression for this plotted curve. This process was repeated to calculate unique read count thresholds for each biological replicate. Read count thresholds were linearly regressed on the total reads for that replicate to conservatively predict thresholds for all samples.

Scaled and filtered reads were normalized by their pseudo gene lengths (theoretical gene length minus number of masked codons) and sequencing depth to give corrected transcripts per million (cTPM). Genes were described as significantly expressed if their cTPM values were among the upper 75th percentile of cTPM values for that sample. For differential expression, genes were described by their fold enrichment between samples if both samples had scaled read counts above their respective read count thresholds. Fold enrichment scores were also used to quantify differential expressions between groups of genes categorized by their ontological function. In cases where only one sample showed read counts above their respective read count threshold, genes were simply described as enriched.

### 5.6 Classification of ORFs

Open reading frames for each gene were characterized using various prediction softwares: clusters of orthologous groups were predicted using EggNOG 4.5 (Huerta-Cepas et al., 2016), subcellular localization was predicted using DeepLoc (Almagro Armenteros et al., 2017), signal sequences were predicted using SignalP 5.0 (Almagro Armenteros et al., 2019), transmembrane domains were predicted using TOPCONS (Tsirigos et al., 2015), and GPI-anchors were predicted using predGPI (Pierleoni et al., 2008). ER-targeting classifications were made for each gene using Ribo-seq data sets from subcellularly-fractionated GS115 Mut^+^ collected during log phase growth in YPD (Alva et al., 2021). These data sets revealed expressions from translating ribosomes in the cytosol and on the membrane of the ER and mitochondria. The log_2_ ratio of cTPM scores for genes in membrane and cytosolic fractions were used to generate membrane enrichment scores. Membrane enrichment scores were used with protein sequence predictions to determine which gene products are translocated into the ER co- and post-translationally. Co-translationally translocated genes had greater than 2-fold membrane enrichment. This classification was broader to include membrane proteins (containing more than two extracytoplasmic transmembrane domains), secreted proteins (containing an N-terminal signal sequence and at most one transmembrane domains near the C-terminus), and proteins without these features that may target the ER using mechanisms involving the 3’UTR. Post-translationally translocated genes show less than 2-fold membrane enrichment and contain a predicted N-terminal signal sequence and less than or equal to one transmembrane domain or a GPI-anchor at the C-terminus. Genes products that met these criteria were filtered to remove those that were predicted to localize to mitochondria.

### 5.7 Rational strain engineering of *K. phaffii* for improved HSA secretion

To validate candidate genes identified by Ribo-seq to rationally improve HSA secretion, *GAL2*, *BGL2*, *YDR134C, SCV12161.1,* and *AOA65896.1* were chosen and knocked out using CRISPR-Cas9 system (**Supplementary Table 4**). The D-227 vector containing *CAS9* and a sgRNA expression cassette (provided by the Love Lab, MIT (Dalvie et al., 2020)) was used for gene knockout. The plasmid was digested with New England BioLabs (NEB) BbVCI restriction enzyme according to the manufacturer’s instructions, and sgRNAs were cloned following our lab’s established protocol (Schwartz and Wheeldon, 2018). Primers for sgRNA cloning were obtained from Integrated DNA Technology (IDT). All cloning was performed with NEBuilder® HiFi DNA Assembly Master Mix (NEB) according to the manufacturer’s instructions. Successful cloning of the sgRNA fragment was confirmed by Sanger sequencing.

After verifying the sgRNA cloning, plasmids were transformed into GS115 Mut^S^ ALB strain using a modified electroporation method (Tafrishi et al., 2024). Transformants were grown in 4 ml of selective media (YPD + 800 μg/ml G418) for two days before being transferred to 2 ml fresh selective media, where they were allowed to grow for an additional two days before plating them on selective plates (YPD, 800 μg/ml G418, and 2% agar). 8 single colonies were randomly picked, the targeting gene was PCR-amplified, and Sanger sequencing was used to verify the successful knockout of the gene (**Supplementary Table 5**).

For HSA quantification, the strain of interest was grown in 200 ml of BMGY until the OD_600_ reached ∼ 6. Next, 2 × 10^10^ cells were transferred to BMMY for HSA induction. Cells were grown in the BMMY media for five days at 30°C and 225 RPM with daily supplementation of 0.5% methanol. After this period, cells were centrifuged at 3000×g for 5 minutes. 100 μl of the supernatant was mixed with 400 μl of Nuclease Free water and was loaded on Amicon^®^ Ultra Centrifugal Filter with a 50 kDa cutoff. Samples were centrifuged at 14000×g for 8 minutes and were washed once again with 500 μl of nuclease free water at 14000×g for 8 minutes. The concentrated supernatant was then recovered and 1 μl of concentrated sample was mixed with 2 μl blue fluorescent protein (BFP), 5 μl of nuclease free water, and 2 ul of 5X SDS Loading buffer. The mixture was heated at 100 ℃ for 5 minutes to denature the proteins and was loaded on SDS-PAGE gels. To quantify the volume of the HSA band, Bio-Rad’s Molecular Imager^®^ ChemiDoc^TM^ XRS System was used. The device’s software was used to identify HSA and BFP bands and to measure their size.

Blue fluorescent protein (BFP) was expressed from plasmid pCRG068 (Supplementary Data) transformed into TOP10 *E. coli* cells. Ni-NTA affinity chromatography was used to purify the polyhistidine-tagged BFP. Briefly, *E. coli* transformants were grown in 100 ml of LB media supplemented with 100 mg/L of ampicillin for 24 hours at 37 ℃ and 225 RPM. Samples were centrifuged, the supernatant was discarded, and cells were distributed in 10 ml of the lysis buffer (50 mM Tris (pH=7.5), 500 mM NaCl). Cells were then sonicated, and the remaining cell pellets were separated from the supernatant using centrifugation. Thermo Scientific disposable plastic columns were packed with 2 ml of Ni-NTA resins according to the manufacturer’s instructions. And the supernatant was loaded on the Ni-NTA resins. After all the supernatant was loaded, the resins were washed three times with the wash buffer (50 mM NaH2PO4 (pH 8.0) and 0.5 M NaCl) and the BFP protein was eluted by gradually adding 2 ml of the elution buffer (3 M Imidazole, 500 mM NaCl, 20 mM Sodium Phosphate Buffer pH = 6.0). To concentrate the purified BFP, Amicon^®^ Ultra Centrifugal Filter with a 10 kDa cutoff was used according to the protocol explained before.

## Author contributions

TA, AT, JC, and IW conceived the project, planned the experiments, and analyzed the data. TA and JC conducted the Ribo-seq experiments, performed and analyzed the NGS data. AT and IW performed genome knockouts and HSA characterization. AT, TA, JC, and IW wrote the manuscript with input from all authors.

## Acknowledgments

Funding from NSF-1951942, -2225878, and -2323984 supported this work. We thank Dr. J. Christopher Love for providing the Cas9 expressing plasmid, D-227.

## Data availability

The Ribo-seq NGS sequencing data generated for this study has been submitted in the NCBI SRA database under accession code PRJNA1222576. Any remaining information can be obtained from the corresponding author upon reasonable request.

