## Supplementary Material for "Ribo-seq guided design of enhanced protein secretion in *Komagataella phaffii*"

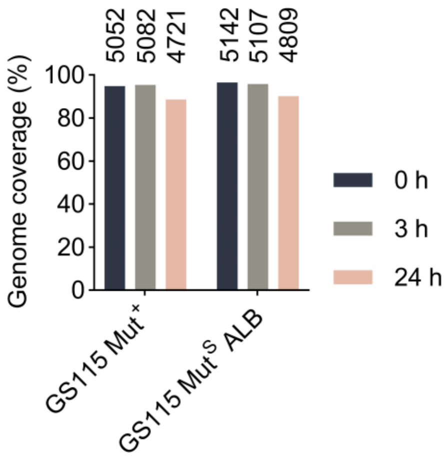


**Supplementary Fig. 1.** Genome-wide coverage of gene expression obtained from Ribo-seq data. The numbers on each bar represent the number of detected genes in the final data.

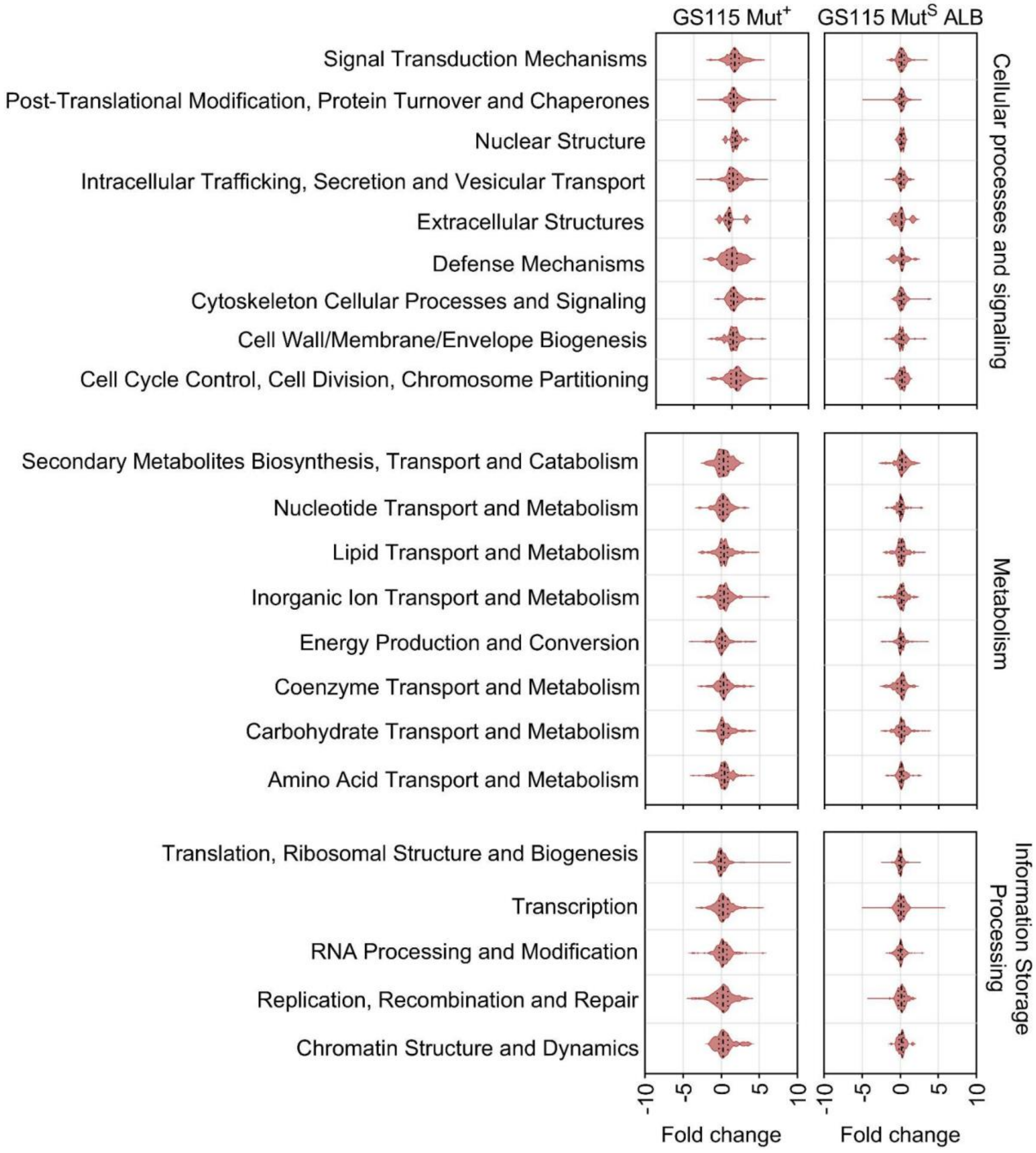


**Supplementary Fig. 2. Fold change production of nascent chains belonging to different ontological categories 3 h after induction.** $\log_2$ (fold change) is calculated as the $\log_2$ ratio of expressed genes 24 hours after methanol induction compared to the expression levels before induction in GS115 Mut$^+$ and GS115 Mut$^S$ ALB and in buffered methanol media (BMMY).

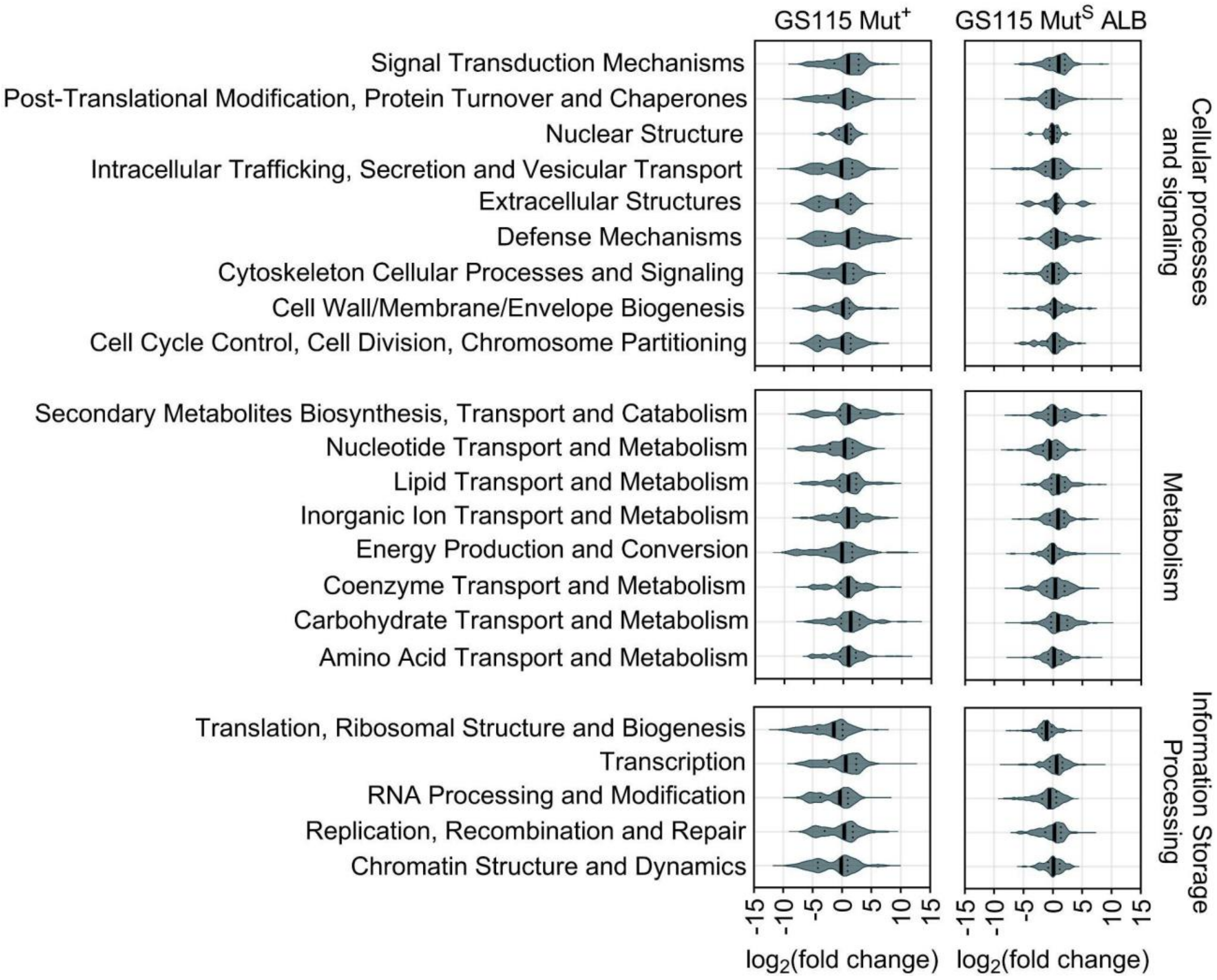


**Supplementary Fig. 3. Fold change production of nascent chains belonging to different ontological categories 24 h after induction**. $\log_2$ (fold change) is calculated as the $\log_2$ ratio of expressed genes 24 hours after methanol induction compared to the expression levels before induction in GS115 $Mut^+$ and GS115 $Mut^S$ ALB and in buffered methanol media (BMMY).. Solid black line shows the median of the plot.

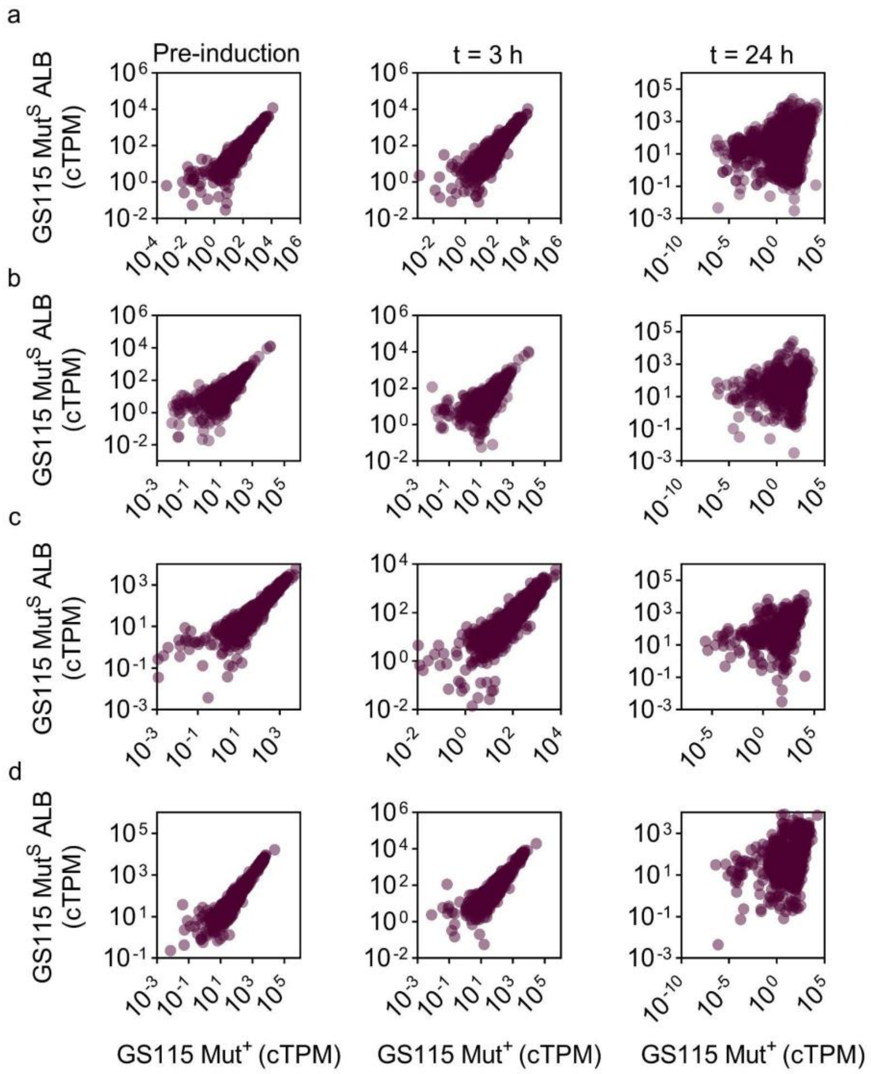


**Supplementary Fig. 4.** Divergence of translational landscape after heterologous expression for ontological categories. **a)** Cell processes and signaling, **b)** Poorly characterized, **c)** Metabolism, and **d)** Information storage and processing

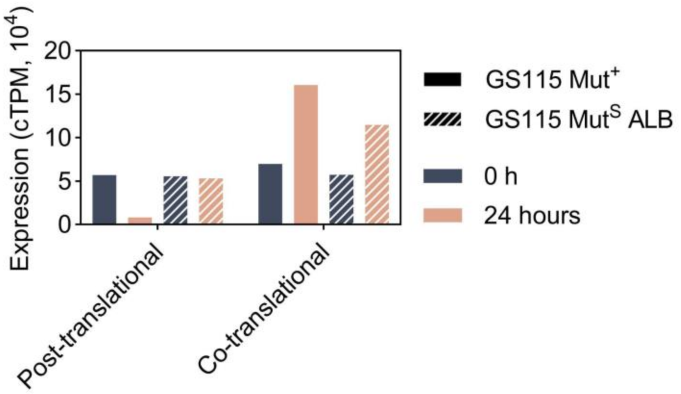


**Supplementary Fig 5. Total nascent chain production translocating into the ER co- and post-translationally.** 56 and 931 protein products are predicted to enter the ER post- and co-translationally.

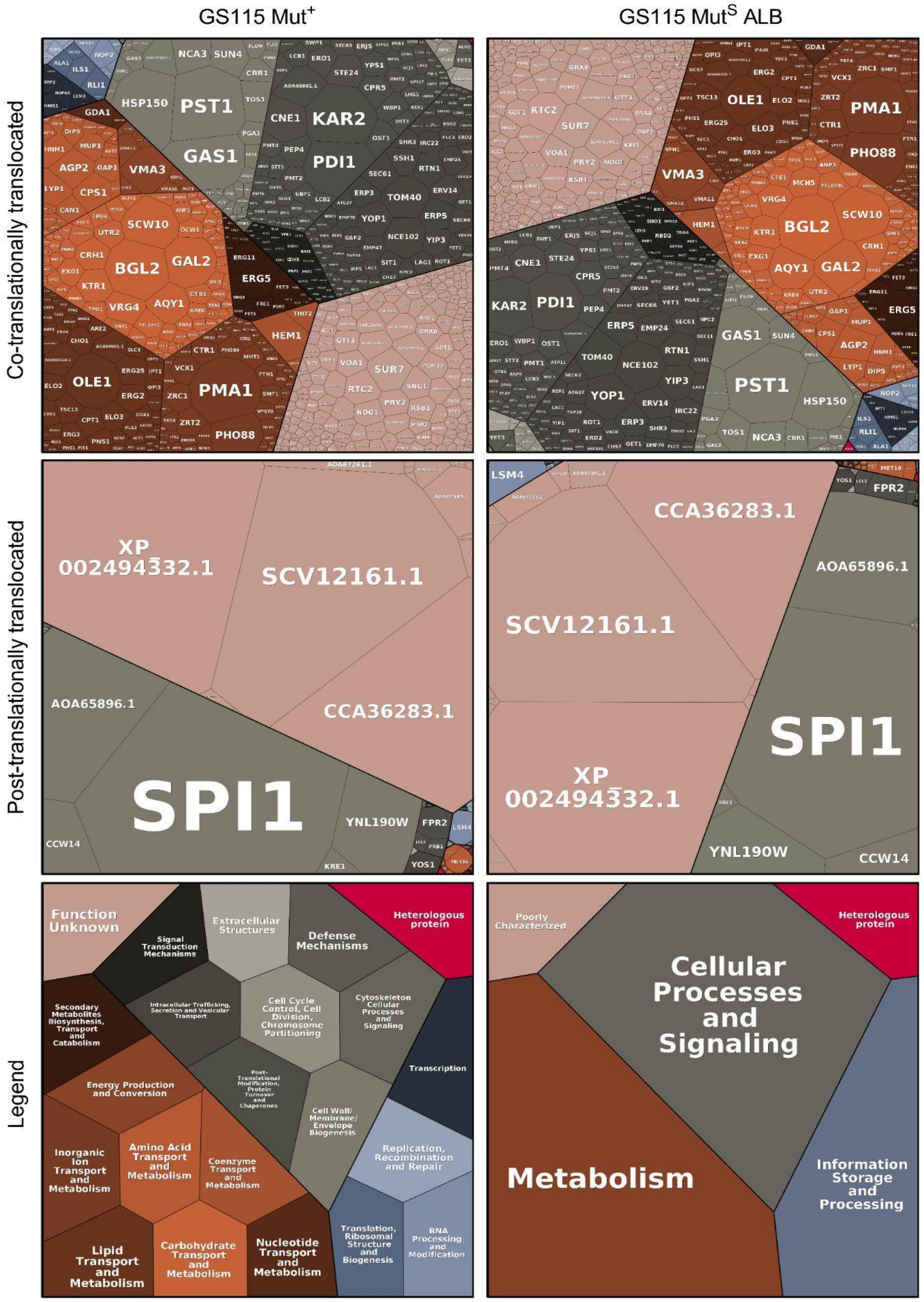


**Supplementary Fig. 6. Co and post-translational flux through the ER for GS115 Mut$^+$ and GS115 Mut$^S$ ALB strains pre-induction.** Non-mitochondrial proteins are predicted to enter the secretory pathway co-translationally if they have greater than $\log_2$ membrane enrichment in YPD studies. Gene products are grouped by ontological function using COG scores predicted by EggNOG v5.0. Cell sizes are calculated using cTPM scores and represent relative quantities of nascent chains produced per gene. Tessellation plots are made using www.bionic-vis.biologie.uni-greifswald.de. [1–3]

**Supplementary Table 1.** Oligos designed for probe-directed degradation

| Probe sequences [a] | Read abundance [b] |
|---|---|
| GTTGGTGCGTCTACGCATCTCCGAC | 10,400,000 |
| CCGTGGGTGAGACGGTCCTAAGGGC | 1,400,000 |
| CATACCCGTGAAAATTTGGTTTATT | 1,000,000 |
| TGTTATTCCCCCGCCCGTACTGACA | 1,000,000 |
| CAAAGAGGGTGATAGCCCCGTGGCA | 760,000 |
| CCTCCGCCCATTCTCAAACTTTAAA | 600,000 |
| AGGGCAGTAAAACCCGAAGAGCGTG | 500,000 |
| CAAAGAGGGTGATAGCCCCGTAGCA | 450,000 |
| TGTGTGGCGAAGACCTGCTTTAGTG | 400,000 |
| GAGTGTTCAAGGCAGTAGTTGAATA | 300,000 |
| ATACAGGGAGGGTGGGGTGAGT | 300,000 |
| CTAGACCCCCTCAGTGGGCCATTTT | 300,000 |
| GTTTAGTTCCATGAGGTAAAGCAAT | 170,000 |
| CGCCAAGGACGTTTTCATTAATCAA | 165,000 |
| ACTCTGGTGGAGGCCCGCAGCGGTT | 130,000 |
| TTATCGACCAACCCAGAACTG | 95,000 |
| CCATATCTAGCAGAAAGCACCGTTT | 86,084 |
| AACGGCGGGAGTAACTATGACTCT | 75,000 |
| AGAAACCTCCAGGCGGGGAGTTTGG | 70,000 |
| ATCGTTGCGAGAGCCAAGAGATCCG | 566 |

[a] Complementary oligonucleotides to Ribo-seq sequences mapped most highly to GS115 rRNA
[b] Ribo-seq reads aligned to GS115 rRNA

**Supplementary Table 2.** Strains used in this study.

| Strain | Genotype | Reference |
| --- | --- | --- |
| E. coli TOP10 | F- mcrA Δ(mrr-hsdRMS-mcrBC) φ80lacZΔM15 ΔlacX74 recA1 araD139 Δ(ara-leu) 7697 galU galK rpsL (StrR) endA1 nupG λ | Thermo Fisher Scientific |
| GS115 Mut$^+$ | *his4* | Invitrogen |
| GS115 Mut$^S$ ALB | *his4, aox1::HSA* | Invitrogen |
| GS115 Mut$^S$ ALB *gal2* | *his4, gal2, aox1::HSA* | This study |
| GS115 Mut$^S$ ALB *aoa65896.1* | *his4, aoa65896.1, aox1::HSA* | This study |
| GS115 Mut$^S$ ALB *bgl2* | *his4, bgl2, aox1::HSA* | This study |
| GS115 Mut$^S$ ALB *scv12161.1* | *his4, scv12161.1, aox1::HSA* | This study |
| GS115 Mut$^S$ ALB *ydr134c* | *his4, ydr134c, aox1::HSA* | This study |
| GS115 Mut$^S$ ALB *gal2 ydr134c* | *his4, gal2, ydr134c, aox1::HSA* | This study |
| GS115 Mut$^S$ ALB *bgl2 ydr134c* | *his4, bgl2, ydr134c, aox1::HSA* | This study |
| GS115 Mut$^S$ ALB *bgl2 gal2* | *his4, gal2, bgl2, aox1::HSA* | This study |
| GS115 Mut$^S$ ALB *scv12161.1 aoa65896.1* | *his4, scv12161.1, aoa65896.1, aox1::HSA* | This study |
| GS115 Mut$^S$ ALB *gal2 aoa65896.1* | *his4, gal2, aoa65896.1, aox1::HSA* | This study |
| GS115 Mut$^S$ ALB *ydr134c aoa65896.1* | *his4, ydr134c, aoa65896.1 , aox1::HSA* | This study |
| GS115 Mut$^S$ ALB *scv12161.1 gal2* | *his4, scv12161.1, gal2, aox1::HSA* | This study |
| GS115 Mut$^S$ ALB *scv12161.1 ydr134c* | *his4, scv12161.1, ydr134c ,aox1::HSA* | This study |
| GS115 Mut$^S$ ALB *gal2 aoa65896.1 ydr134c* | *his4, gal2, aoa65896.1, ydr134c, aox1::HSA* | This study |
| GS115 Mut$^S$ ALB *gal2 aoa65896.1 ydr134c* | *his4, gal2, aoa65896.1, ydr134c, aox1::HSA* | This study |
| GS115 Mut$^S$ ALB *gal2 bgl2 ydr134c* | *his4, gal2, bgl2, ydr134c, aox1::HSA* | This study |
| GS115 Mut$^S$ ALB *gal2 aoa65896.1 ydr134c scv12161.1* | *his4, gal2, aoa65896.1, ydr134c, scv12161.1, aox1::HSA* | This study |
| GS115 Mut$^S$ ALB *gal2 aoa65896.1 ydr134c bgl2* | *his4, gal2, aoa65896.1, ydr134c, bgl2, aox1::HSA* | This study |
| GS115 Mut$^S$ ALB *gal2 aoa65896.1 ydr134c bgl2 scv12161.1* | *his4, gal2, aoa65896.1, ydr134c, bgl2, scv12161.1 , aox1::HSA* | This study |

**Supplementary Table 3.** Barcode sequences used for demultiplexing NGS data.

| Barcode sequence | Sample |
|---|---|
| 'NNNNNATCGTAGATCGGAAGAGCACACGTCTGAA' | Total fraction; t = 0 h |
| 'NNNNNCGTAAAGATCGGAAGAGCACACGTCTGAA | Total fraction; t = 3 h |
| 'NNNNNGATCAAGATCGGAAGAGCACACGTCTGAA' | Total fraction; t = 24 h |

**Supplementary Table 4.** sgRNAs used in this study.

| sgRNA target | sgRNA sequence |
|---|---|
| *AOA65896.1* | ACAAGAGGTGATAGTCAGAA |
| *YDR134C* | GAACTCTCCAGAATCAGCAA |
| *GAL2* | GACATCCCAGTCAAACCCAA |
| *BGL2* | GGAATAAGCTCTAATAGCAA |
| *SCV12161.1* | TTCGTTTTGAGCTTGCACAA |

**Supplementary Table 5.** Primers used in this study.

| Primer Name | Primer sequence |
|---|---|
| AOA65896.1.FOR | TTCCTCAACTCACTGTTTCAGTTTATTCCAAC |
| AOA65896.1.REV | GTGAGAGCTGGTCTTAGCTGGAG |
| BGL2.FOR | ATCTGAAGCTGGCAAGTCGTC |
| BGL2.REV | GATCTTTAATCTTAAAACACTGGCTGCG |
| GAL2.FOR | TAATATGAGTTCAACAGATATCCAAGGTGATCAAG |
| GAL2.REV | AAGGTAATACGTTTCACCGTTAAACTGT |
| SCV12161.1.FOR | CCACAAAATTTCAGCGAGCAACAG |
| SCV12161.1.REV | AGTCCTCACCTACAGCCAACT |
| YDR134C.FOR | ATAATGCTAACCAAGGTTATTTCACTCGC |
| YDR134C.REV | AGTGTAAGAAACACATTCGGGGT |
